# Chronic exposure of mosquito larvae to pesticide residues endangers a new generation of agrochemicals repurposed for malaria prevention

**DOI:** 10.1101/2023.04.18.537423

**Authors:** Marilene M. Ambadiang, Caroline Fouet, Fred A. Ashu, Véronique Penlap-Beng, Colince Kamdem

**Affiliations:** Centre for Research in Infectious Diseases (CRID), P.O. Box 13591, Yaoundé, 9 Cameroon; Department of Biochemistry, Faculty of Science, University of Yaoundé 1, P.O. Box 11 812, Yaoundé, Cameroon; Department of Biological Sciences, The University of Texas at El Paso, 500 W. University Ave. El Paso, TX 79968

**Keywords:** pesticide, neonicotinoids, *Anopheles*, malaria, urbanization

## Abstract

Agrochemicals have been successfully repurposed to control mosquitoes worldwide, but pesticides used in agriculture challenge their effectiveness by contaminating surface waters and helping larval populations develop resistance. Thus, knowledge of the lethal and sublethal effects of residual pesticide exposure on mosquitoes is critical for selecting effective insecticides. Here we implemented a new experimental approach to predict the efficacy of agricultural pesticides newly repurposed for malaria vector control. We mimicked insecticide resistance selection as it occurs in contaminated aquatic habitats by rearing field-collected mosquito larvae in water containing a dose of insecticide capable of killing individuals from a susceptible strain within 24 h. We then simultaneously monitored short-term lethal toxicity within 24 h and sublethal effects for 7 days. We found that due to chronic exposure to agricultural pesticides, some mosquito populations are currently pre-adapt to resist neonicotinoids if those were used in vector control. Larvae collected from rural and agricultural areas where neonicotinoid formulations are intensively used for insect pest management were able to survive, grow, pupate and emerge in water containing a lethal dose of acetamiprid, imidacloprid or clothianidin. These results emphasize the importance of addressing exposure of larval populations to formulations applied in agriculture prior to using agrochemicals against malaria vectors.

## 1. Introduction

Prevention through chemical control of vector populations has lead to a drastic reduction in malaria burden in sub-Saharan Africa over the last two decades (World Health Organization, 2022a). The prevention strategy relies heavily on repurposing agrochemicals, which provides a rapid and cost-effective approach to identify new active ingredients against vector populations (Hemingway, 2017; Hoppé et al., 2016). In order to effectively control *Anopheles* mosquito populations that have become resistant to available insecticides, dozens of agrochemicals have recently been tested against adult populations, and some promising candidates have been identified (Hoppé et al., 2016; Lees et al., 2019; N’Guessan et al., 2007; Oxborough et al., 2010; Portwood et al., 2022). However, concomitant use of the same class of agrochemicals for crop protection can provide sufficient selection pressure to engineer resistant mosquito populations before any application of the pesticide in public health (Georghiou, 1972; Georghiou et al., 1973; Lines, 1988). For example, prior to the scale up of pyrethroid-impregnated bed nets across the continent, resistance driven by crop protection spraying was already detected in some *Anopheles* populations in West Africa (Curtis et al., 1998). There is a concern history may repeat itself with the new generation of agrochemicals that are being adopted for malaria vector control (World Health Organization, 2023).

Africa’s pesticide use for agriculture stood at nearly 108 thousand metric tons in 2019 (Kramer, 2022). Lax regulations allow the use of hundreds of formulations for agricultural pest management in some African countries (Akpesse, 2018; Katambo, 2018; Ngamo Tinkeu, 2018; Okolle et al., 2022; Urio et al., 2022). A large part of pesticide application occurs in small farms held by non-professional operators that are not trained and not supervised for safe and effective use of chemicals. As a result, too frequent and untargeted application provides suitable conditions for chronic exposure of mosquito larvae to pesticide residues. In tropical regions, rain and human activities create ponds in farmlands, which become ideal breeding sites for the most dangerous malaria mosquito species such as *Anopheles gambiae* and *An. coluzzii* in agricultural areas (Antonio-Nkondjio et al., 2011; Mattah et al., 2017; Zogo et al., 2019). When these breeding sites are contaminated with pesticides, chronic residual exposure pre-adapts larval populations to resist synthetic chemicals (Lines, 1988). The consequence of this adaptation is a decrease in efficacy of agrochemicals, which in turn undermines their suitability for mosquito control.

Neonicotinoids are among the most widely used pesticides in agriculture worldwide (Matsuda et al., 2020; Simon-delso et al., 2015). These chemicals are highly water-soluble and environmentally persistent, and thus may leach into surface waters and become a risk for aquatic insects (Morrissey et al., 2015; Ramadevi et al., 2022; Schaafsma et al., 2015). Two new formulations of clothianidin, a neonicotinoid, were recently approved for indoor residual spraying targeting malaria vectors, but have yet to be applied in endemic countries (Agossa et al., 2018; Kweka et al., 2018; Uragayala et al., 2018; World Health Organization, 2023). Additionally, clothianidin is normally not registered for crop protection in most African countries (Katambo, 2018; Ngamo Tinkeu, 2018). So far, agricultural spraying of other registered neonicotinoids such as acetamiprid, imidacloprid, thiamethoxam and thiacloprid remain the main source of selection pressure that could drive resistance to clothianidin in *Anopheles* mosquitoes through cross-resistance mechanisms. This hypothesis has yet to be supported by empirical evidence, but observational studies have described suboptimal levels of neonicotinoid susceptibility in adults of at least two *Anopheles* species, populations from agricultural areas being the most tolerant (Ashu et al., 2023a; Fouet et al., 2020; Mouhamadou et al., 2019). Importantly, there is evidence that neonicotinoids used for crop protection can confer cross-resistance to clothianidin, raising concerns about the efficacy the new agrochemicals repurposed for malaria prevention (Ashu et al., 2023a, 2023b; Fouet et al., 2020).

Large African cities are the venue of dynamic coevolution between humans and disease vectors, which has sometimes reveled textbook examples of rapid adaptation of mosquitoes to synthetic chemicals (Antonio-Nkondjio et al., 2015; Kamdem et al., 2017, 2012b). In Yaoundé the capital of Cameroon, agricultural activities associated with intensive use of pesticides are pervasive in suburban and rural settings in the outskirts of the city. Mixtures containing neonicotinoids such as acetamiprid, imidacloprid, and thiamethoxam, as well as pyrethroids and fungicides are frequently sprayed on diverse crops, creating ideal conditions for contamination of surface waters and resistance selection (Ngamo Tinkeu, 2018; Okolle et al., 2022). By contrast, the center of the city provides an island where surface waters are less exposed to pesticide contamination and aquatic species are therefore less likely to develop resistance. Indeed, a survey of susceptibility in adult mosquitoes using standard testing procedures confirmed this prediction. *An. coluzzii*, the only species found in densely urbanized areas of Yaoundé remains susceptible to neonicotinoids while its sibling species *An. gambiae*, which occurs in the countryside, has developed resistance (Ashu et al., 2023a, 2023b; Fouet et al., 2020). This geographic area thus provides a unique setting to dissect the environmental drivers of neonicotinoid resistance in *Anopheles* mosquitoes.

The objective of the present study was to assess the interplay between residual pesticide exposure and the development of resistance to new insecticides in malaria vectors. It has been known for nearly a half century that exposure to agricultural pesticides can lead to the spread of resistance to public health insecticides in *Anopheles* mosquitoes (Georghiou, 1972; Georghiou et al., 1973). However, we still lack a holistic approach that could allow to simultaneously deciphering the lethal (mortality) and sub-lethal effects (survival, growth, emergence, reproduction, swimming behavior, mobility etc.) which contribute to resistance selection in agricultural settings. To fill this gap, we implemented a bioassay protocol where we mimicked pesticide exposure in the wild by rearing filed-collected *Anopheles* larvae in water containing field-realistic doses of chemicals. Using larval populations collected from urban and rural areas of Yaoundé, we compared mortality within 24 h as well as some life table parameters in order to infer the role of neonicotinoid contaminants in resistance selection. Our findings indicated that resistance selection in *An. gambiae* is due to chronic exposure to neonicotinoids in rural and agricultural settings and translates into low mortality but also high survival, growth, pupation and emergence in lethal concentrations of pesticides. The present study highlights concerns regarding the efficacy of some agrochemicals repurposed for malaria vector control, especially active ingredients that are concomitantly used in agricultural pest management.

## 2. Material and methods

### 2.1 Study sites

The study was carried out in Yaoundé, the capital of Cameroon, and some of its neighboring rural areas. Approval to conduct a study in the Center region (N°: 1-140/L/MINSANTE/SG/RDPH-Ce), ethical clearance (N°: 1-141/CRERSH/2020) and research permit (N°: 000133/MINRESI/B00/C00/C10/C13) were granted by the ministry of public health and the ministry of scientific research and innovation of Cameroon. We surveyed four sites which included a farming area located in the suburbs (Nkolondom, 3°56’43’’ N, 11°3’01’’ E), two densely urbanized neighborhoods (Etoa Meki, 3°52’53’’ N, 11°31’40’’ E and Combattant, 3°52’53’’ N, 11°31’40’’ E) and another suburban site (Nkolnkoumou, 3°52’29’’ N, 11°23’2’’ E) (Figure 1). The average distance between sites was 4-5 km. The most abundant malaria vector populations in this area belong to the species complex *An. gambiae sensu lato (s*.*l*.) (Antonio-Nkondjio et al., 2011; Kamdem et al., 2012a, 2012b). Previous surveys have revealed that the nominal species *An. gambiae sensu stricto* (hereafter *An. gambiae*) is the only member of the complex present in the agricultural site Nkolondom while its sibling species *An. coluzzii* prefers the most urbanized areas of Yaoundé (Bamou et al., 2019; Kamdem et al., 2012b; Tene-Fossog et al., 2022). Larvae from the semirural neighborhood, Nkolnkoumou, are a mixture of ∼ 80% *An. gambiae* and 20% *An. coluzzii* (Ashu et al., 2023a, 2023b; Fouet et al., 2020). In rural and urban settings, larvae thrive in rain-dependent puddles, which may contain high concentrations of organic pollutants in some densely urbanized areas of the city (Antonio-Nkondjio et al., 2011). In the agricultural site Nkolondom, *An. gambiae* larvae occur in waters that accumulate between furrows and ridges in farms due to rain and/or irrigation and are likely contaminated with pesticide residues (Figure 1).

**Figure 1.**
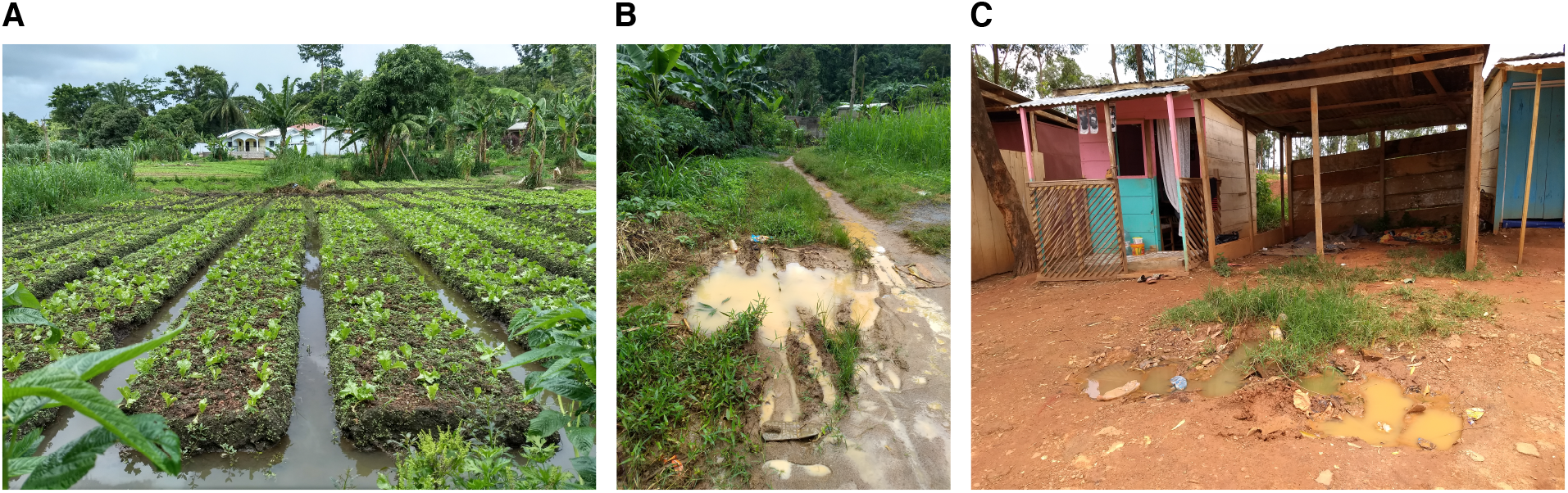
Examples of breeding sites where wild mosquito larvae were sampled. *An. gambiae* larvae were collected from furrows between ridges in the agricultural site (A) or from puddles created by human activities in Nkolnkoumou (suburban area) (B). *An. coluzzii* larvae were sampled from human-made puddles in densely urbanized settings (C).

### 2.2 Larval collection and insecticides

Typical *Anopheles gambiae s*.*l*. breeding sites were inspected during rainy seasons in 2022 to collect larvae using the standard dipping method (Service, 1993). Larvae were transported in plastic containers to the insectary where they were immediately tested in a controlled room (27°C, 80% relative humidity, light:dark = 12:12 h).

Three neonicotinoids were tested in this study: in addition to acetamiprid and imidacloprid that are used in local farming activities, we also tested clothianidin, which remains an exotic insecticide in Cameroon. Deltamethrin, a first-generation malaria-control insecticide, was used to compare the efficacy of clothianidin. The following commercial formulations were tested: acetamiprid (Aceplant 40EC, 40g/L, emulsifiable concentrate, JACO, Yaoundé, Cameroon), imidacloprid (Plantima 30SC, 30g/L, concentrated suspension, JACO, Yaoundé, Cameroon), clothianidin (Pestanal™, analytical standard, Sigma-Aldrich) and deltamethrin (Decis 25EC, 25 g/L, emulsifiable concentrate, Bayer Cropscience SL, Valencia, Spain). Stock solutions were prepared by diluting the formulation in absolute ethanol or distilled water.

### 2.3 Larval bioassays

In order to assess the lethal and sublethal effects of each insecticide, we reared larvae collected from the field in water containing a dose of the insecticide that was lethal to larvae of a susceptible strain and we measured mortality as well as some life history parameters. The mosquito life cycle comprises four larval stages that precede pupation and emergence of a male or female adult (Service, 1993). The four stages include the 1^st^ (L1), 2^nd^ (L2), 3^rd^ (L3) and 4^th^ (L4) instars and can last between one and two weeks depending on the mosquito species, the feeding regime and the environmental conditions. We monitored mortality and life table parameters starting from 3^rd^ instars until emergence. 3^rd^ instar larvae of *Anopheles* are abundant in the wild, are easy to identify morphologically and typically complete transformation into L4, pupation and emergence in approximately 7 days.

The doses tested for each insecticide were determined as follows: we exposed 3^rd^ instar larvae of the susceptible strain *An. gambiae* Kisumu to increasing concentrations of the insecticide, starting from 0.001 mg/L and we selected the lowest concentration at which 100% of individuals died within 24 h. Batches of 25 larvae were placed in 500 mL plastic trays filled with 200 mL borehole water containing the desired concentration of insecticide and covered with a net. Four replicates were tested for each insecticide dose in addition to a control without insecticide (i.e. containing only water). The lowest lethal concentrations were 0.035 mg/L clothianidin, 0.075 mg/L imidacloprid, 0.15 mg/L acetamiprid and 1.5 mg/L deltamethrin. According to studies conducted worldwide, the concentrations of neonicotinoids tested fell in the upper limits of average values detected in contaminated waters and soils (Morrissey et al., 2015; Ramadevi et al., 2022; Schaafsma et al., 2015). However, in some agricultural areas, these concentrations can be 100 times higher (Ramadevi et al., 2022).

Larvae were collected from the field in the morning and brought to the insectary. 3^rd^ instars were sorted immediately and rinse in a tray containing borehole before being transferred into test trays. Four replicates of 25 larvae were tested in 500 mL plastic trays that were filled with 200 mL borehole water containing the above-determined lethal dose of the insecticide. Water was not changed throughout the experiment, and 10 mg of TetraMin® fish food was added to each tray daily. Every 24 h, the number of L3, L4, pupae and adults was counted in each tray. Larvae were considered dead if they were unable to move when touched with a dropper. Dead larvae were removed from the test containers and were not replaced. Adults were also removed using a mouth aspirator. Survival, growth, pupation and emergence were assessed daily for 7 days, which was sufficient for 3^rd^ instar larvae to reach the adult stage.

### 2.4 Data analysis

The lethal endpoint of insecticide exposure was assessed using the mortality rate at 24 h. The sublethal effects of the different insecticides were addressed using survival probability as well as L4 transformation rate, pupation rate and emergence rate. L4 transformation rate was defined as the percentage of L3 larvae transformed into L4. Pupation rate represented the percentage of L3 that reached the pupa stage and emergence rate the percentage of L3 that became adults. Line plots of mean and standard error, computed with the packages *plyr* and *ggplot2* in R V. 4.2.2, were used to compare L4 transformation rate, pupation rate and emergence rate at 24 h intervals. The ratio between the number of dead larvae and the initial number of larvae was calculated every 24 h and provided an estimate of survival probability. Kaplan Meier survival curves were plotted using the packages *ggplot 2* and *ggfortify* in R (R Core Team, 2016). Larvae reaching the adult stage before the end of the test were treated as censored data. Confidence intervals were computed for the four replicates, and a log-rank test was used to determine if survival was significantly different between treatments.

## 3. Results

### 3.1 Lethal toxicity in 24 h

We first tested a gradient of insecticide doses on the laboratory strain *An. gambiae* Kisumu, which is susceptible to common classes of insecticides used in mosquito control such as pyrethroids, organophosphates, carbamates and organochlorines. The objective was to determine a dose yielding 100% mortality in 24 h. We then used this dose as a discriminating concentration to study the lethal and sublethal effects of neonicotinoids on field collected *Anophleles* larvae. We compared the efficacy of neonicotinoids to that of deltamethrin. As opposed to the susceptible strain, 100% mortality was not achieved with field populations after 24 h (Figure 2). Mortality rates were very low in larvae of both *An. gambiae* and *An. coluzzii* reared in water containing deltamethrin. Pyrethroid resistance is widespread in anopheline mosquitoes, and parallel application of deltamethrin in malaria prevention and in agricultural pest management has drastically reduced its efficacy against disease vectors. 24-h mortality in water containing deltamethrin was typically below 30% and was not significantly different between the four tested populations (Wilcoxon rank sum test, p>0.05). In contrast with deltamethrin, susceptibility to neonicotinoids varied between species and geographic areas. Consistent with findings obtained using adult bioassays (Ashu et al., 2023a; Fouet et al., 2020), *An. coluzzii* larvae collected from the two urban neighborhoods, Combattant and Etoa meki, were more susceptible, with mortality rates between 70-80%, except clothianidin for which only about 50% of larvae were killed within 24 h. The species present in suburban and rural areas of Yaoundé, *An. gambiae*, has developed strong resistance to neonicotinoids. Particularly, larvae collected from the Nkolondom farm where pesticide spraying has selected for resistance to acetamiprid and imidacloprid and conferred cross-resistance to clothianidin (Ashu et al., 2023a; Fouet et al., 2020), displayed less than 10% mortality in water containing a neonicotinoid. Mortality rate of larvae collected from the agricultural site was <10% in water containing the new malaria-control chemical, clothianidin, or the first-generation insecticide, deltamethrin, which suggested that both active ingredients currently have similar efficacy against some *An. gambiae* populations.

**Figure 2.**
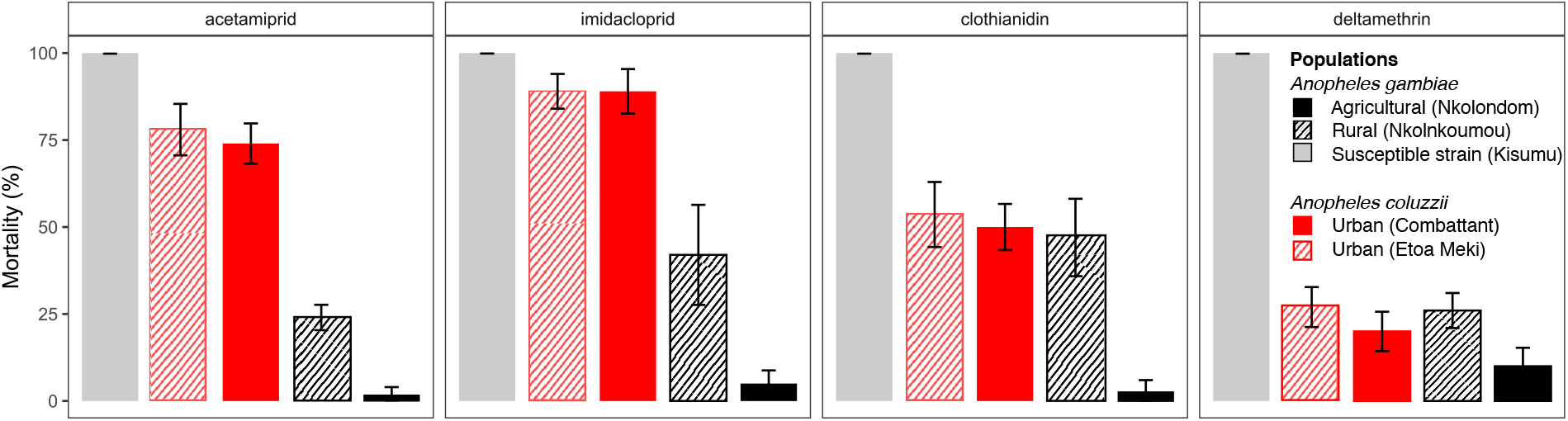
24-h mortality of mosquito larvae reared in water containing a lethal dose of agrochemical. 100% of larvae of the susceptible strain *An. gambiae* Kisumu were killed within 24 h. Wild populations showed varying levels of tolerance to four insecticides. Mortality values were lowest in larvae collected from the agricultural area (Nkolondom). Error bars represent the standard error of the mean.

### 3.2 L4 transformation rate

Some neonicotinoid insecticides act slowly and are also known to have a wide range of sublethal effects in addition to acute toxicity on non-target species (Morrissey et al., 2015; Schuhmann et al., 2022). Thus, beside lethal toxicity within the first 24 h of the experiment, we used life table analysis to monitor some of the sublethal effects of neonicotinoids on *Anopheles* larvae, starting with L4 transformation rate. The three neonicotinoids tested had little effect on the ability of *An. gambiae* L3 larvae to become L4 (Figure 3A). The highest transformation rate was observed in populations from the agricultural site, Nkolondom: 90 ± 3% in water containing acetamiprid followed by 75 ± 9% in imidacloprid in two days and 60 ± 3% in clothianidin after 3 days. Similarly, in Nkolnkoumou, a semi-rural site harboring ∼ 80% *An. gambiae*, transformation rates varied from 60 ± 6% for acetamiprid, to 35 ± 16% and 23 ± 7% for imidacloprid and clothianidin, respectively after 2 to 4 days. Conversely, even if based on 24-h mortality, *An. coluzzii* larvae showed a degree of tolerance, they were strongly inhibited in water containing neonicotinoids leading to very low transformation rates between 0 and 10% in both Etoa Meki and Combattant populations. In control tests, transformation rates were 100% after 2 to 3 days.

**Figure 3.**
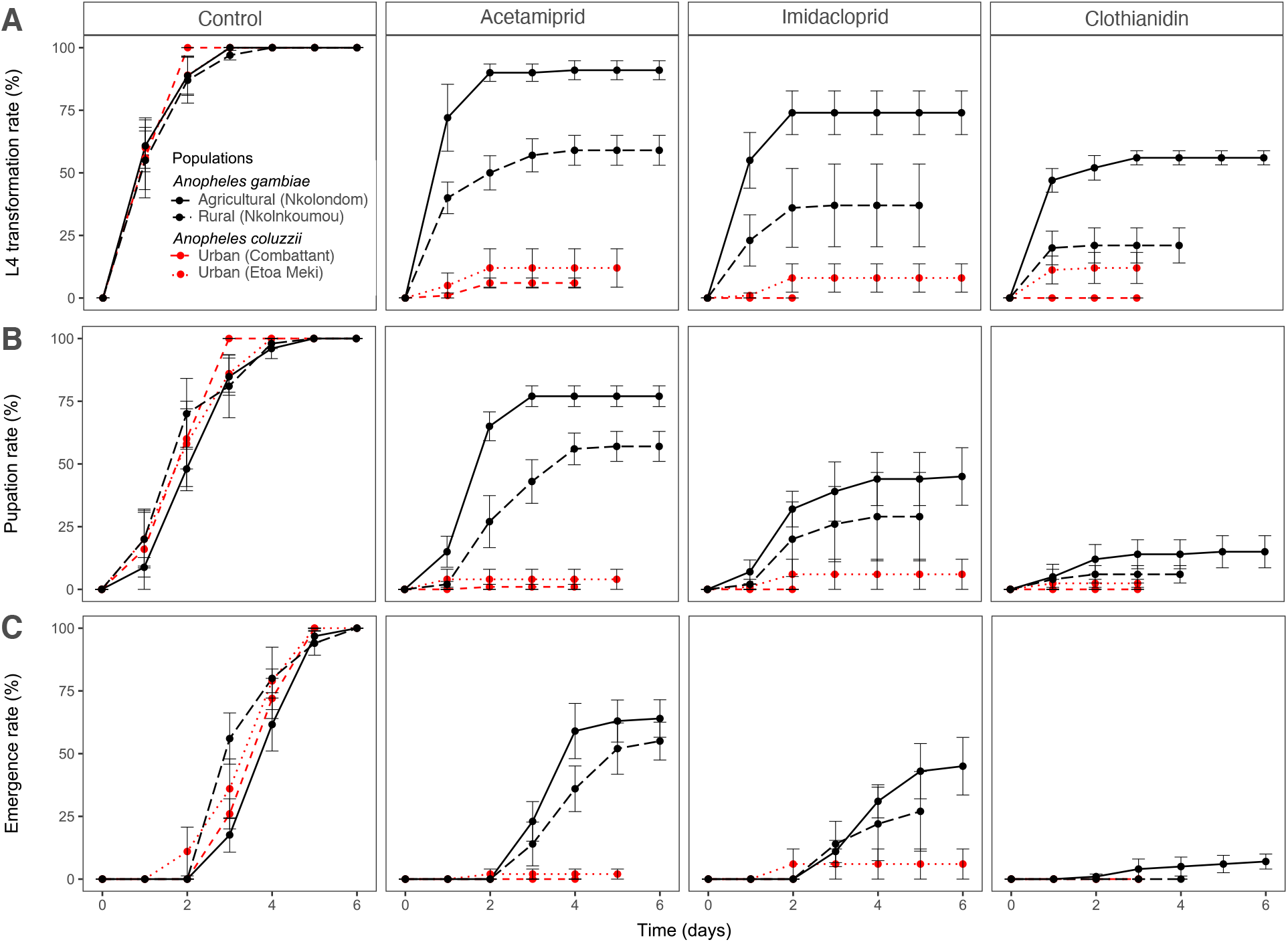
Sublethal effects of three neonicotinoids on life table parameters of *Anopheles* larvae. Rate of transformation of 3^rd^ instars (L3) into 4^th^ instars (L4) (A), pupation rate (B) and emergence rate (C). Larvae that survived 24-h lethal toxicity were monitored for 7 days under standard laboratory conditions. Controls were larvae reared in water without insecticide. Vertical bars represent the standard error of the mean.

### 3.3 Pupation rate

Pupation rate was consistent with L4 transformation and highlighted the low susceptibility of agricultural and semi-rural populations of *An. gambiae* to neonicotinoids (Figure 3B). The agricultural population of *An. gambiae* (Nkolondom) had the highest pupation rate, 77.5 ± 4% in water containing acetamiprid, and 45 ± 11% in imidacloprid reflecting their tolerance to the both neonicotinoid pesticides that are sprayed in local farms. The exotic insecticide, clothianidin, at 0.035 mg/L had the most important inhibitory power among neonicotinoids, but 15 to 20% of *An. gambiae* larvae were able to achieve pupation. *Anopheles coluzzii* larvae had low pupation rates with less than 5% pupae obtained in all the neonicotinoids tested. Notably, none of the *An. coluzzii* populations were able to pupate in clothianidin.100% pupae were obtained in all the control tests after 72 h.

### 3.4 Emergence rate

100% of *An. gambiae* and *An. coluzzii* pupae emerged into adults in control water without insecticide between the first and the sixth day in standard laboratory conditions. The total emergence rate of *An. coluzzii* larvae was 0%, 2% and 4% in water containing clothianidin, acetamiprid or imidacloprid, respectively after 7 days (Figure 3C). Meanwhile, at least half of *An. gambiae* L3 larvae tested on average emerged between 2-7 days in acetamiprid and imidacloprid. Only *An. gambiae* larvae from the agricultural site were able to emerge in clothianidin albeit at lower rate (7 ± 3%) compared with the other neonicotinoids. However, emergence rate was also very low in deltamethrin, around 10%, indicating that for any insecticide, the ability to inhibit emergence can be compatible with strong resistance in adult mosquitoes and loss of efficacy of the insecticide. In addition to lethal toxicity in 24 h, emergence rate revealed another similarity between clothianidin, a new insecticide and deltamethrin whose effectiveness in malaria vector control has become very limited.

### 3.5 Survival probability

Rearing field-collected larvae in water containing an agricultural pesticide affected their survival probability as revealed by Kaplan Meier survival curves (Figure 4). Larvae that survived lethal toxicity within the first 24 h of the experiment were monitored for seven days. Although *An. coluzzii* showed potential to resist the 24-h lethal effect of some of the agricultural pesticides, larvae of this species had low survival probability in neonicotinoids. By contrast, *An. gambiae* populations from agricultural and semi-rural sites displayed high survival potential in water containing lethal doses of neonicotinoids. A log-rank test revealed that regardless of the neonicotinoid that was added to water, larvae collected from Nkolondom were super-resistant and had significantly higher survival probability compared with any of the three other populations tested (p < 0.05). 100% survival was obtained in all control experiments without insecticide.

**Figure 4.**
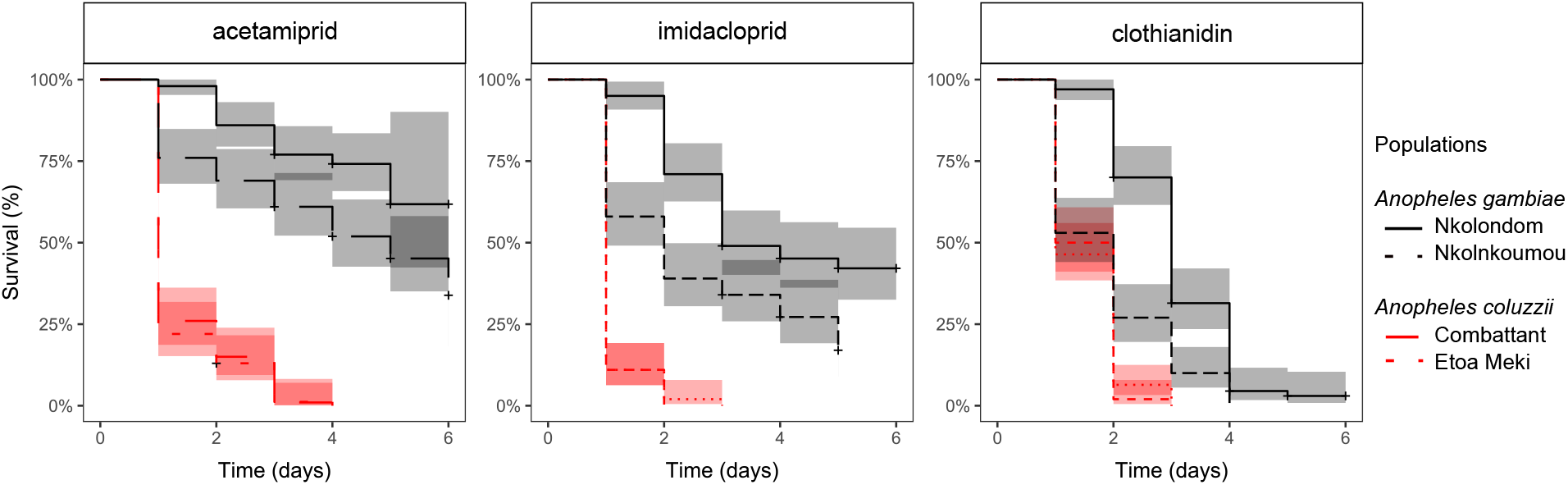
Kaplan Meier survival curves (lines) with 95% confidence intervals (colored bands) of 3^rd^ instar larvae reared in water containing a lethal dose of neonicotinoid. Survival probabilities of four wild populations were compared under controlled laboratory conditions. (+) indicates emergence that occurred before the end of the experiment and was treated as censored data.

## 4. Discussion

Here we have used an innovative approach to simultaneously assess the lethal and sublethal effects of pesticide exposure in mosquito larvae. We found that monitoring some sublethal effects via persistent exposure to chemicals in a controlled aquatic environment provides critical information on the level of tolerance of larval populations. Such information could be complementary to routine quantification of mortality in adults via standard bioassays (CDC, 2012; World Health Organization, 2022b), and combining both approaches may offer a more robust framework to better evaluate pre-existing levels of adaptation to agrochemicals in malaria mosquitoes. We conducted a laboratory experiment mimicking the selection process, which is promoting the emergence of *Anopheles* populations with increased tolerance to neonicotinoids. While analyzing mortality and life table in wild larvae reared in artificial media containing a lethal dose of the active ingredient, we observed gradients of tolerance reflecting past exposure to neonicotinoids in nature. Control tests supported the validity of our experimental approach as 3^rd^ instar larvae unexposed to insecticides showed standard survival, growth and emergence in controlled insectary conditions.

Previous studies have suggested that resistance to neonicotinoids observed in anopheline populations from some agricultural areas could be due to chronic exposure of larvae to sublethal doses of agricultural pesticides (Ashu et al., 2023a, 2023b; Fouet et al., 2020; Mouhamadou et al., 2019). Our findings provide the first experimental support to this hypothesis. Indeed, we found that *An. gambiae* larvae had exceptional fitness in field-realistic concentrations of acetamiprid and imidacloprid, two neonicotinoids whose formulations are extensively used in farming activities in Cameroon (Fouet et al., 2020). As anopheline populations have not yet been exposed to clothianidin in sub-Saharan Africa, this exotic neonicotinoid strongly inhibited larval emergence in both *An. gambiae* and *An. coluzzii*. However, likely due to cross-resistance selected by intensive exposure to acetamiprid and imidacloprid, larvae from the agricultural site, Nkolondom, were able to survive in water contaminated with clothianidin, albeit at lower frequency.

Exposure of non-target insects to sublethal doses of neonicotinoids affect functions such as motility, behavior, growth, fecundity and survival (Morrissey et al., 2015; Řezáč et al., 2021; Shepherd et al., 2021). Our study reveals that notwithstanding these sublethal effects whose magnitude remains unknown, *An. gambiae* populations have evolved adaptation to chronic exposure to neonicotinoid residues in contaminated aquatic habitats. *An. gambiae* larvae from rural areas around Yaoundé exhibited high survivorship, L4 transformation, pupation and emergence rate in water containing neonicotinoids contrary to urban *An. coluzzii* mosquitoes whose growth and survival were inhibited. This result corroborates susceptibility profiles of adult populations: *An. coluzzii* from Yaoundé is more susceptible to neonicotinoids while *An. gambiae* is resistant to at least four different active ingredients (Ashu et al., 2023a, 2023b; Fouet et al., 2020). Larval populations whose adults were resistant to an agrochemical (i.e. mortality against the discriminating dose < 90%) typically displayed less than 50% mortality in 24 h and more than 5% emergence in 6 days in water containing a lethal dose of the active ingredient. Testing a larger number of chemicals on diverse *Anopheles* species and populations will provide more robust guidelines on the interpretation of mortality, growth and emergence of larvae reared in pesticide-laced water. However, based on the current study and complimentary evidence from adults testing (Ashu et al., 2023a, 2023b; Fouet et al., 2020), we could establish that any agrochemical yielding less that 50% mortality after 24 h and more than 5% emergence within 6 days is likely to have limited efficacy against *Anopheles* mosquitoes.

The first evidence of insecticide resistance driven by agricultural pesticide application dates back to the 1960s. Based on observations made in El Salvador, it was clearly established that aerial spraying of carbamates and organophosphates led to the development of broad-spectrum resistance to both insecticide classes in the vector *Anopheles albimanus* (Georghiou, 1972; Georghiou et al., 1973). Since then, a number of observational studies have argued that intensive use of crop protection chemicals is a major driver of resistance to public health insecticides in anopheline mosquitoes (Chouaïbou et al., 2016; Diabate et al., 2002; Müller et al., 2008; Nwane et al., 2009; Urio et al., 2022; Yadouleton et al., 2009). However, in the majority of cases, it was difficult to determine the guilty part between agricultural chemicals and malaria vector control insecticides. In a review of evidence, Lines (1988) (Lines, 1988) proposed a set of conditions that should be met for a crop protection chemical to be the source of insecticide resistance selection in malaria vectors. Appearance of resistance in mosquitoes to a particular chemical should occur before it has been used for mosquito control; there should be correlation in space and time, e.g. higher levels of resistance in areas with agricultural spraying and seasonality of resistance overlapping with spraying periods; the cross-resistance spectrum should show that the chemicals used in agriculture may confer cross-resistance to public health insecticides and that both adults and larvae can be killed by the agricultural pesticides. Agricultural application of neonicotinoids fits all of the criteria in Sub-Saharan Africa and provides the best example of human activities accidentally engineering insecticide resistance in mosquitoes.

In our study, both lethal and sublethal effects point to the existence of levels of resistance that could be a major obstacle to using neonicotinoids in malaria mosquito control. We noted that lethal toxicity in 24 h, pupation rate and emergence rate were very similar in water containing either deltamethrin or clothianidin. This finding suggested that, just like deltamethrin, the efficacy of clothianidin in *An. gambiae* might already be substantially reduced. In line with this prediction, it has been shown that neonicotinoid resistant mosquitoes from Nkolondom were significantly less susceptible to SumiShield® 50WG, a formulation of clothianidin recently approved for indoor residual spraying (Fouet et al., 2020). Moreover, a laboratory experiment recently demonstrated that exposure of *Anopheles* larvae to sublethal concentrations of a mixture containing several herbicides, pesticides and fungicides resulted in ∼ 2.5 increase in tolerance to clothianidin and Fludora® Fusion, a formulation combining clothianidin and deltamethrin (Zoh et al., 2022). These studies and ours highlight the vulnerability of some agrochemicals repurposed for malaria vector control, as resistance driven by agricultural activities could become a major thread to their efficacy.

Although our findings are based on only two *Anopheles* species sampled from a relatively small geographic area, these results confirm concerns about the efficacy of neonicotinoids against malaria vectors (Ashu et al., 2023a; Fouet et al., 2020; Hoppé et al., 2016; Mouhamadou et al., 2019; Oxborough et al., 2019). Repurposing agrochemicals has thus far provided a rapid mechanism to identifying new candidate insecticides for malaria prevention. However, the case of neonicotinoids emphasizes the crucial role of evaluating prior exposure and cross-resistance in areas where mosquito larval populations are likely to be in contact with agricultural pesticides.

## Author Contributions

MA: Conceptualization, Formal analysis, Investigation, Methodology, Writing – original draft; CF: Conceptualization, Formal analysis, Investigation, Methodology; FA: Investigation, Methodology; VP-B, Resources, Supervision. CK: Conceptualization, Formal analysis, Funding acquisition, Investigation, Project administration, Writing – review & editing.

## Informed Consent Statement

Not applicable.

## Data Availability Statement

The data for this study have been presented within this article.

